# Loss of lamin-B1 and defective nuclear morphology are hallmarks of astrocyte senescence *in vitro* and in the aging human hippocampus

**DOI:** 10.1101/2021.04.27.440997

**Authors:** Isadora Matias, Luan Pereira Diniz, Isabella Vivarini Damico, Laís da Silva Neves, Ana Paula Bergamo Araujo, Gabriele Vargas, Renata E. P. Leite, Claudia K. Suemoto, Ricardo Nitrini, Wilson Jacob-Filho, Lea T. Grinberg, Elly M. Hol, Jinte Middeldorp, Flávia Carvalho Alcantara Gomes

## Abstract

The increase in senescent cells in tissues, including the brain, is a general feature of normal aging and age-related pathologies. Senescent cells exhibit a specific phenotype, which includes an altered nuclear morphology and transcriptomic changes. Astrocytes undergo senescence *in vitro* and in age-associated neurodegenerative diseases, but little is known about whether this process also occurs in physiological aging. Here, we investigated astrocyte senescence *in vitro*, in old mouse brains and in *post-mortem* human brain tissue of elderly. We identified a significant loss of lamin-B1, a major component of the nuclear lamina, as a hallmark of senescent astrocytes. We showed a severe reduction of lamin-B1 in the dentate gyrus of aged mice, including in hippocampal astrocytes, and in the granular cell layer of the hippocampus of *post-mortem* human tissue from non-demented elderly. Interestingly, the lamin-B1 reduction was associated with nuclear deformations, represented by an increased incidence of invaginated nuclei and loss of nuclear circularity in senescent astrocytes *in vitro* and in the aging human hippocampus. In conclusion, our findings show that reduction of lamin-B1 is a conserved hallmark of astrocyte aging, as well as shed light on significant defects in nuclear lamina structure, which may impact astrocyte function during human aging.

## INTRODUCTION

Aging is characterized by a progressive change in the physiology of brain cells that may contribute to cognitive deficits, leading ultimately to dementia and impairment of quality of life. The unique complexity of the human brain, as well as the increasing longevity of humans, represent additional challenges to prevent and postpone aging effects.

It is estimated that by 2050 the number of people aged 60 and older is double from now, reaching nearly 2.1 billion worldwide (Nations, 2017). In this context, a substantial increase in the incidence of age-associated diseases, such as cancer, diabetes, and neurodegenerative diseases is expected for the next few years (Hou et al., 2019). However, the mechanisms underlying the transition from a healthy fully functional brain to an elderly increasingly dysfunctional brain are not well understood.

Among the causal factors of aging, cellular senescence is an important tumor suppression mechanism, but it also leads to a harmful cellular phenotype which affects tissue homeostasis and regeneration (Campisi, 2005; Lopez-Otin et al., 2013). Several studies have reported age-related accumulation of senescent cells in humans, non-human primates and rodents (Herbig et al., 2006; Wang et al., 2009; Yousefzadeh et al., 2020). In the aging brain, neural stem cells, neurons and glial cells display several features of cellular senescence and have been implicated in the pathogenesis of age-related neurodegenerative diseases, such as Alzheimer’s Disease (AD) and Parkinson’s Disease (PD) (Cohen and Torres, 2019; Martínez-Cué and Rueda, 2020). However, so far little is known about the impact of glial cell senescence on normal brain aging.

Astrocytes form a large and heterogeneous glial cell population in the central nervous system (CNS), playing several vital roles in brain homeostasis and physiology (Verkhratsky and Nedergaard, 2018). In contrast, astrocyte dysfunction has been associated with many brain disorders, such as AD and PD (Diniz et al., 2019; Matias et al., 2019; Osborn et al., 2016), in which senescent astrocytes accumulate and are linked to neurodegeneration and cognitive and motor impairments in mouse models (Bhat et al., 2012; Bussian et al., 2018; Chinta et al., 2018). Nevertheless, despite the amount of evidence pointing to the involvement of astrocyte senescence in age-related pathologies, the phenotypic and functional changes of senescent astrocytes in the context of physiological brain aging remain to be elucidated. This is at least partially due to the lack of senescence-associated biomarkers for glial cells. Moreover, most of our knowledge of astrocyte biology and brain aging comes from rodent models. A growing number of studies shows the unique complexity of human astrocytes (Barbar et al., 2020; Oberheim et al., 2009) and thus it is important to study whether the mechanisms underlying rodent brain aging also apply to the human brain.

The senescent state usually requires a coordinated activation of well-known molecular changes, which identify senescent cells *in vitro* and *in vivo* (Sharpless and Sherr, 2015). Moreover, senescent cells undergo important morphological changes, including altered nuclear size and shape (Mehta et al., 2007), as well as reduced levels of lamin-B1 (Freund et al., 2012).

Lamin-B1, together with the other nuclear lamins, constitute the major components of the nuclear lamina, where they form a dense filamentous meshwork essential for maintaining the structural and functional integrity of the nucleus (de Leeuw et al., 2018). Mutations in lamin genes cause several devastating human diseases, collectively known as laminopathies, among which are premature aging syndromes (Schreiber and Kennedy, 2013). Interestingly, nuclear deformations and reduced levels of B-type lamins are present in *post-mortem* human brain tissue of AD and Frontotemporal Dementia (FTD) patients (Frost et al., 2016; Paonessa et al., 2019). Besides, lamin-B1 loss was found specifically within astrocytes in PD brain tissue, suggesting the contribution of senescent astrocytes to PD pathology (Chinta et al., 2018). Still, it remains to be elucidated whether lamin-B1 levels and nuclear morphology are altered in astrocytes during normal brain aging.

Here, we investigated whether lamin-B1 loss and changes in nuclear morphology accompany astrocyte senescence *in vitro* and in the mouse and human aging brain. Altogether, our findings suggest that lamin-B1 loss and defective nuclear morphology are hallmarks of astrocyte senescence *in vitro* and human hippocampal aging.

## RESULTS

### Age-related loss of lamin-B1 in the mouse hippocampus

Loss of lamin-B1 has been described as a senescence-associated biomarker in some tissues *in vivo* (Saito et al., 2019; Wang et al., 2017; Yue et al., 2019), although it remains less known whether this also applies to the aged brain. To address this question, we first examined with immunofluorescence the levels of lamin-B1 in the hippocampal dentate gyrus (granular cell layer and molecular layer) of C57Bl/6 mice at 2-3 months and 18-24 months of age. We observed a steady reduction of more than 75% of lamin-B1 intensity in aged mice compared with young ones **(Figure 1 A-G)**. Interestingly, in young mice, lamin-B1 intensity was higher at the subgranular zone and in a few cells of the molecular layer **(Figure 1 C-C’)**. In contrast, there was a huge decline of lamin-B1 throughout the molecular and granular cell layers of the dentate gyrus in aged mice **(Figure 1 F-F’)**.

**Figure 1.**
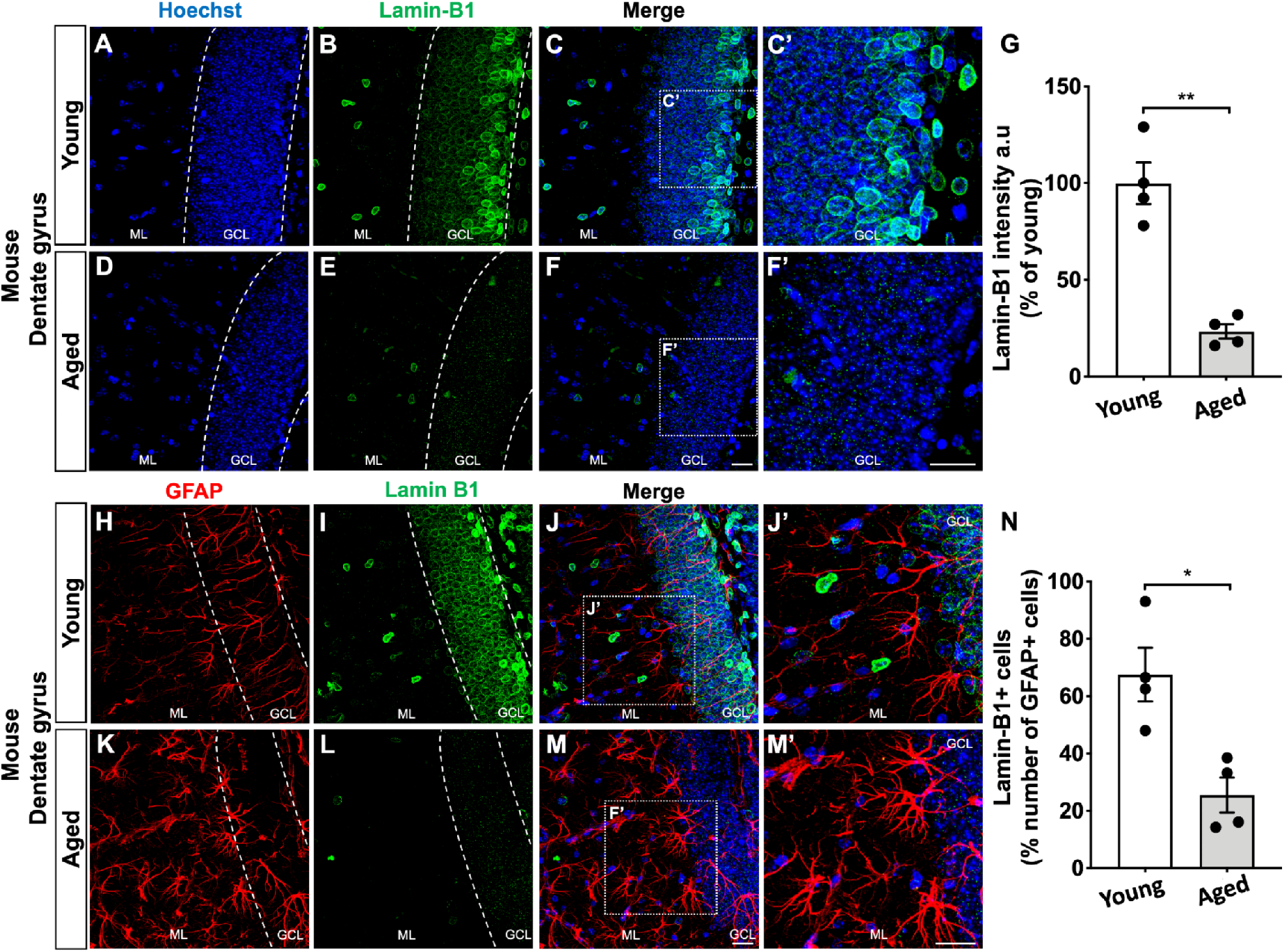
Age-related loss of lamin-B1 in the mouse hippocampus. (A-G) Densitometric analysis of lamin-B1 staining in the mouse hippocampal dentate gyrus, including the molecular layer (ML) and granular cell layer (GCL), revealed a global reduction of lamin-B1 intensity in aged mice compared with young mice (p=0.0033). (H-N) Decreased proportion of lamin-B1+ cells in relation to the total number of GFAP+ cells in the dentate gyrus of aged mice compared with young ones (p=0.0126). Significance was determined using Unpaired t test with Welch’s correction. Error bars represent ± SEM. Individual data points are plotted and represent individual animals (*n*=4 animals per experimental group). Scale bars, 20 µm.

To further address the subcellular decrease of lamin-B1 in the aged hippocampus, we evaluated lamin-B1 staining in GFAP+ cells in young and aged mice. While we found ∼65% of lamin-B1+/GFAP+ cells at hippocampal dentate gyrus of young mice, aged mice showed 25% of lamin-B1+/GFAP+ cells **(Figure 1 H-N)**.

Altogether, these results point to an age-related reduction of lamin-B1 in the mouse hippocampal dentate gyrus, which is at least partially due to a decline of expression of this molecule in astrocytes.

### Lamin-B1 reduction is a hallmark of senescent astrocytes *in vitro*

Lamin-B1 downregulation has been described as a senescence-associated biomarker in different *in vitro* models for cellular senescence, including in replicative, oncogene, and DNA damage-induced senescence models (Freund et al., 2012; Limbad et al., 2020). However, less is known about the regulation of lamin-B1 in aged glial cell cultures.

We thus sought to investigate lamin-B1 as an aging biomarker of astrocytes as suggested by hippocampal mouse data. To do that, we first established a new *in vitro* model for astrocyte senescence by culturing mouse cortical astrocytes for 9-10 days *in vitro* (DIV, control group) or 30-35 DIV (senescent group) followed by evaluation of several features of cellular senescence **(Figure 2 A)**.

**Figure 2.**
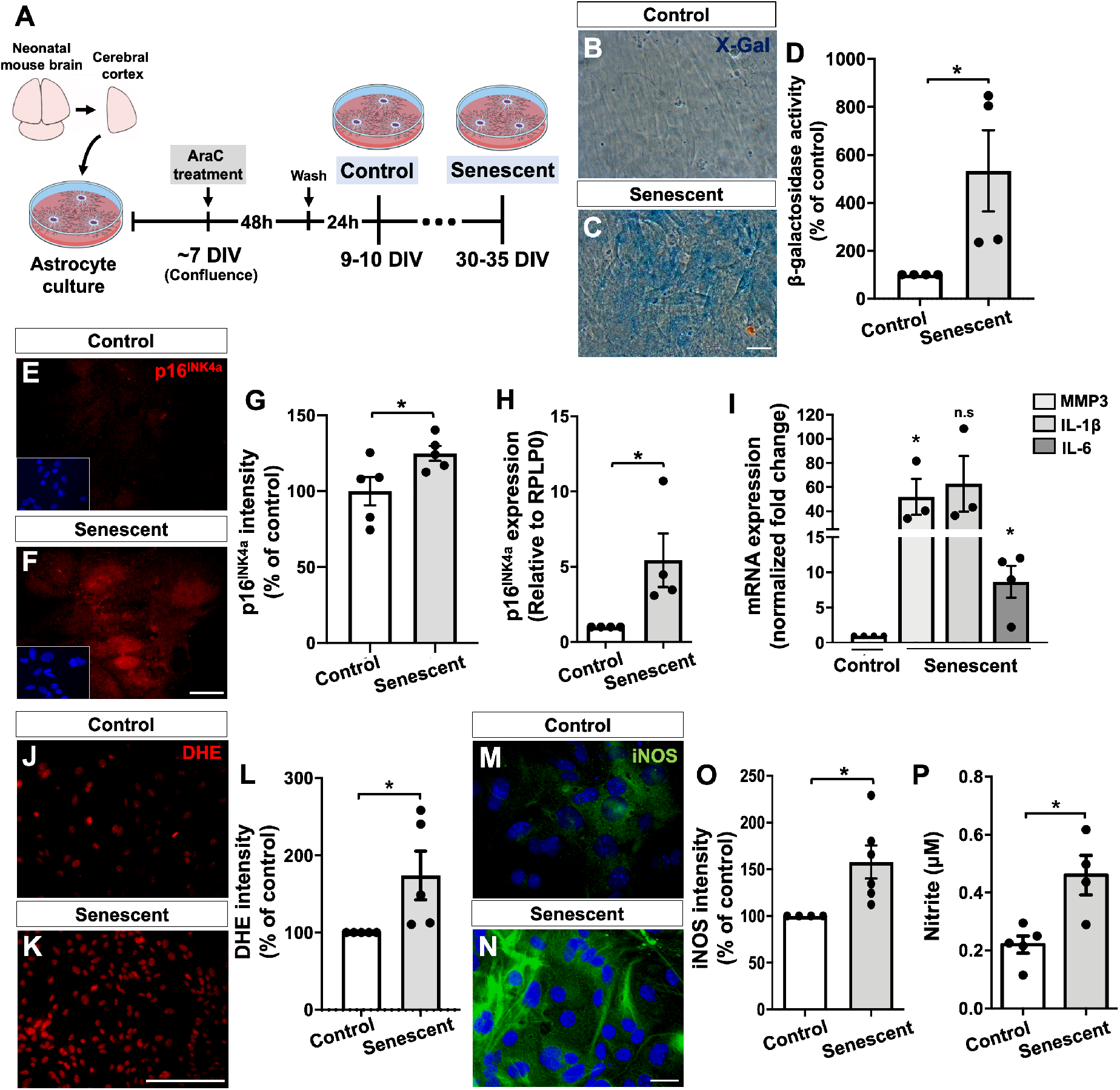
Characterization of a new *in vitro* model for astrocyte senescence. (A) Primary murine astrocyte cultures were maintained for 9-10 DIV (control group) or 30-35 DIV (senescent group) and analyzed for several senescence-associated biomarkers. (B-D) Astrocytes cultured for 30-35 DIV showed increased SA-β-galactosidase activity compared with control cultures (p=0.0426). (E-G) Higher immunostaining for p16^INK4a^ in senescent astrocytes compared with control cultures (p=0.0459). (H) Senescent astrocytes showed increased expression of p16^INK4a^ (p=0.0471). (I) Expression levels of MMP3 and IL-6 were increased in senescent astrocyte cultures compared with control ones (p=0.0267 and p=0.0142, respectively). (J-L) Increased intensity of DHE labelling (level of ROS) in senescent astrocytes compared with control cultures (p=0.0467). (M-O) Higher immunostaining for iNOS in senescent astrocytes (p=0.0311). (P) Nitrite production was elevated in senescent astrocytes (p=0.0100). Significance was determined using Unpaired t test. Error bars represent ± SEM. Individual data points are plotted and represent individual cultures (*n*=3-6 cultures per experimental group). Scale bars, 20 µm.

We first analyzed the astrocyte cultures for two classical senescence-associated biomarkers: SA-β-Gal and the expression of the tumor suppression protein, p16^INK4a^ (Sharpless and Sherr, 2015). Accordingly, we observed increased activity of SA-β-Gal **(Figure 2 B-D)**, as well as increased intensity and expression of p16^INK4a^ in astrocytes cultured for 30-35 DIV compared with control astrocytes **(Figure 2 E-H)**.

Senescent cells are known to express a senescence-associated secretory phenotype (SASP), linked to increased secretion of several inflammatory mediators, matrix metalloproteinases, and reactive oxygen and nitrogen species (Coppé et al., 2010). In line with this, we observed an upregulation of matrix metalloproteinase 3 (MMP3) and IL-6 mRNA in senescent astrocytes compared with the control group **(Figure 2 I)**. Moreover, senescent astrocytes also showed increased levels of reactive oxygen species **(Figure 2 J-L)**, as well as higher intensity of inducible nitric oxide synthase (iNOS, **Figure 2 M-O**) and extracellular nitrite (NO_2_^-^), a stable metabolite of nitric oxide **(Figure 2 P)**.

Immunostaining and Western blotting assays revealed a robust reduction in lamin-B1 intensity **(Figure 3 A-C)** and its protein level **(Figure 3 D)**, of approximately, 50% and 30%, respectively, in senescent astrocytes compared with control cultures.

**Figure 3.**
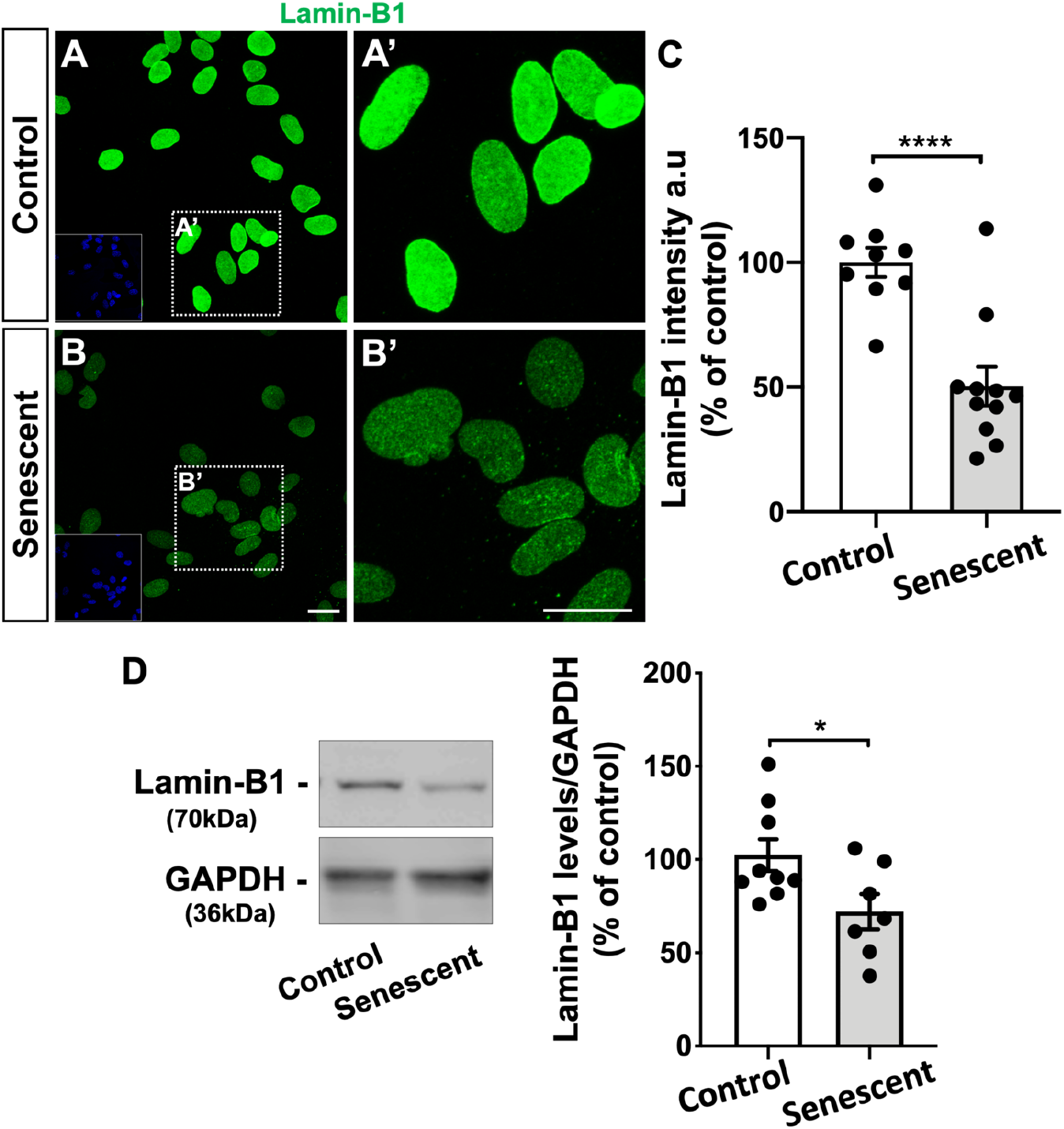
Lamin-B1 reduction is a hallmark of senescent astrocytes *in vitro*. (A-C) Reduced immunostaining for lamin-B1 in senescent astrocytes compared with the control group (p<0.0001). *n*=9 and 11 independent astrocyte cultures for control and senescent groups, respectively. Scale bars, 20 µm. (D) Lamin-B1 protein levels were diminished in senescent astrocytes in comparison with the control group (p=0.0329). *n*=9 and 7 independent cultures for control and senescent groups, respectively. Significance was determined using Unpaired t test with Welch’s correction. Error bars represent ± SEM. Individual data points are plotted and represent individual cultures.

Taken together, these results indicate first, that mouse astrocytes cultured for 30-35 DIV recapitulate key features of the senescent phenotype, and validate their use as an alternative *in vitro* model for astrocyte senescence. In addition, these results corroborate previous reports on the lamin-B1 decline in cellular senescence models and provide a new hallmark for astrocyte senescence.

### Nuclear deformations are associated with lamin-B1 loss in cultured senescent astrocytes

It has been shown that lamin concentration within the nucleus is essential in regulating nuclear lamina assembly and therefore in maintaining nuclear structure and function (Guo et al., 2014; Paonessa et al., 2019). We thus sought to investigate the impact of lamin-B1 downregulation in astrocyte nuclear morphology.

We first analyzed the nuclear circularity based on lamin-B1 staining, which has a maximum value of 1 and diminishes as the nuclear shape becomes increasingly convoluted. We observed a significant reduction in nuclear circularity in senescent astrocytes compared with control ones **(Figure 4 A-E)**, which had a positive correlation with the reduced intensity of lamin-B1 in senescent cells **(Figure 4 F)**.

**Figure 4.**
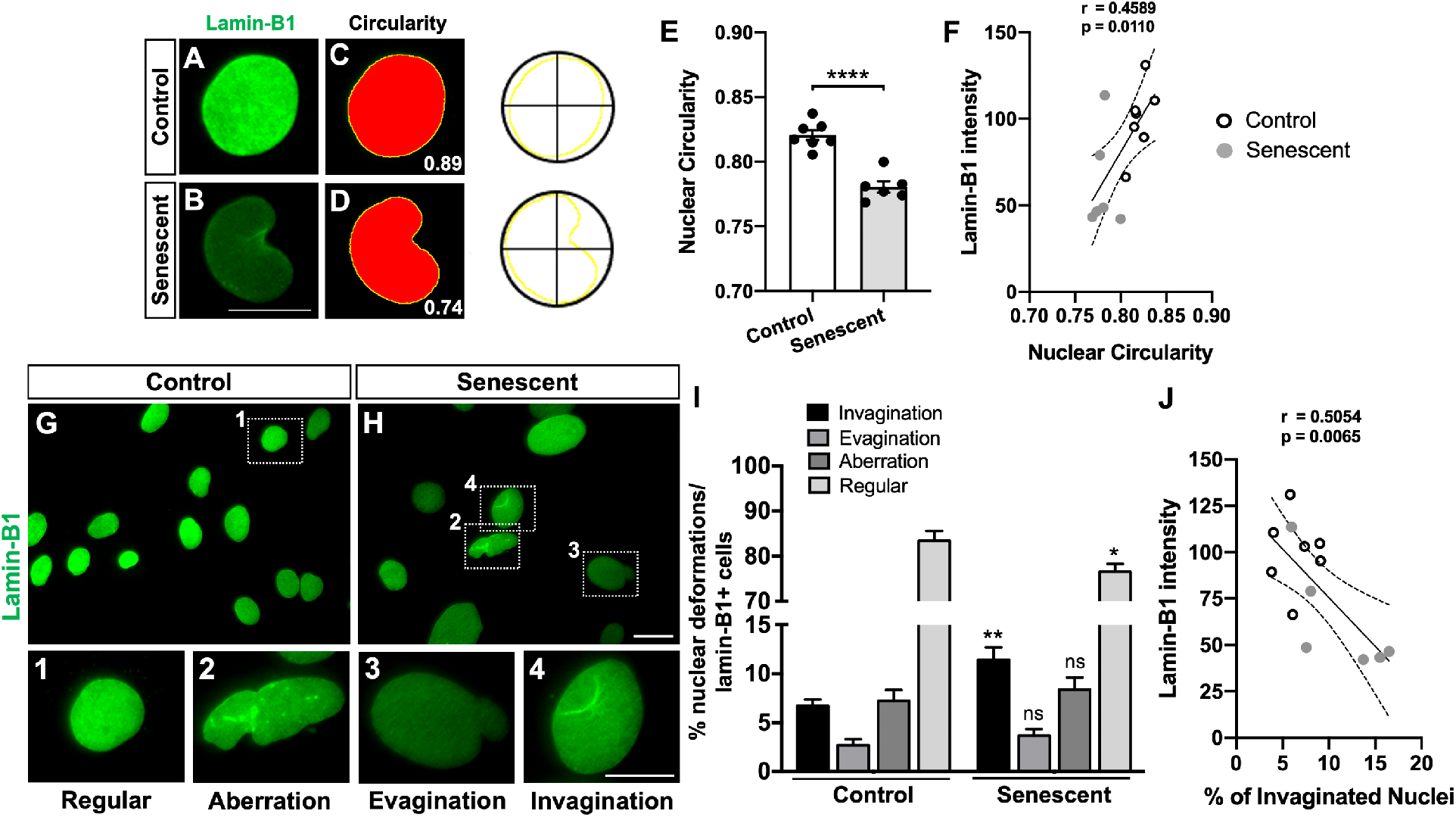
Nuclear deformations are associated with lamin-B1 loss in senescent astrocytes. (A-D) Nuclear circularity analysis is based on the area and perimeter of the nucleus. Circularity has a maximum value of 1 and diminishes as the nuclear shape becomes increasingly convoluted, as observed in senescent cells (B, D). (E) Senescent astrocytes displayed a reduced nuclear circularity compared with the control group (p<0.0001). *n*=7 and 6 cultures for control and senescent groups, respectively. (F) A positive correlation was observed between lamin-B1 intensity and the nuclear circularity value (r=0.4589; p=0.0110). *n*=7 and 6 independent cultures for control and senescent groups, respectively. (G-H) Distinct nuclear morphological profiles were evaluated, such as regular (1), aberration (2), evagination (3) and invagination (4) in control and senescent cultured astrocytes, based on lamin-B1 staining. Scale bars, 20 µm in (H) and 10 µm in (4). (I) Senescent astrocyte cultures showed a significant increased incidence of invaginated nuclei (p=0.0054) and a decreased proportion of regular nuclei (p=0.0263) compared with control cultures. *n*=9 and 10 independent cultures for control and senescent groups, respectively. (J) A negative correlation was observed between lamin-B1 intensity and the incidence of invaginated nuclei (r=0.5054; p=0.0065). *n*=7 and 6 independent cultures for control and senescent groups, respectively. Significance was determined using Unpaired t test with Welch’s correction. Linear regression opting to show 95% confidence bands of the best-fit line. Error bars represent ± SEM. Individual data points are plotted and represent individual cultures. Control cultures are represented by white dots and senescent cultures by gray dots in (F) and (J).

We next asked whether alterations of the nuclear circularity could be associated with other types of nuclear lamina abnormalities. To do so, we initially performed three-dimensional reconstruction of lamin-B1+ nuclei of astrocytes, based on z-stack fluorescence microscopy, to classify the types of nuclear deformations present in control and senescent astrocyte cultures. The nuclear morphological classification was divided into three types: invagination, when nuclei showed one clear lamin-B1 invagination; evagination, when nuclei exhibited one clear lamin-B1 protrusion from the nuclear lamina; and aberration, when nuclei presented a combination of more than one invagination, evagination or additional nuclear abnormalities. Regular nuclei were considered those without any of these deformations.

Therefore, in order to optimize the analysis, we used conventional fluorescence microscopy to image and quantify the number of nuclear deformations in control and senescent astrocyte cultures. We observed a reduced proportion of regular nuclei in senescent astrocyte cultures compared with control astrocytes (**Figure 4 G-I)**. This was mainly due to the increased incidence of nuclear invaginations in these cells **(Figure 4 G-I)**. The proportion of nuclear evaginations and aberrations was similar between control and senescent astrocytes **(Figure 4 G-I)**. Interestingly, the incidence of nuclear invaginations negatively correlated with lamin-B1 intensity in astrocytes **(Figure 4 J)**.

Therefore, these results strongly indicate that astrocyte senescence *in vitro* is associated with defects in nuclear morphology, represented by nuclear invaginations and reduced nuclear circularity, which correlate with reduced levels of lamin-B1 in these cells.

### Lamin-B1 reduction in the human dentate gyrus upon aging

Whether the mechanisms underlying rodent brain aging apply to the human brain is still a matter of investigation. Thus, we further evaluated whether lamin-B1 downregulation also occurs in the human brain during normal aging. To do so, we gathered *post-mortem* human hippocampal tissue from two independent brain banks, both composed of non-demented controls separated into two age groups: middle-aged and elderly donors. Clinic-pathological information of all donors is present in **Supplementary Table 1**.

Densitometric analysis of lamin-B1 immunostaining revealed an overall reduction of approximately 15% in lamin-B1 intensity at the granular cell layer of the hippocampal dentate gyrus in the elderly compared with middle-aged donors **(Figure 5)**. It is noteworthy that this hippocampal layer is mainly composed by granule neuron cells, and at its inner boundery reside neural stem/progenitor cells and glial cells, especially astrocytes (Casse et al., 2018). Therefore, these results indicate that human aging is accompanied by a downregulation of lamin-B1 in the granular cell layer of the dentate gyrus.

**Figure 5.**
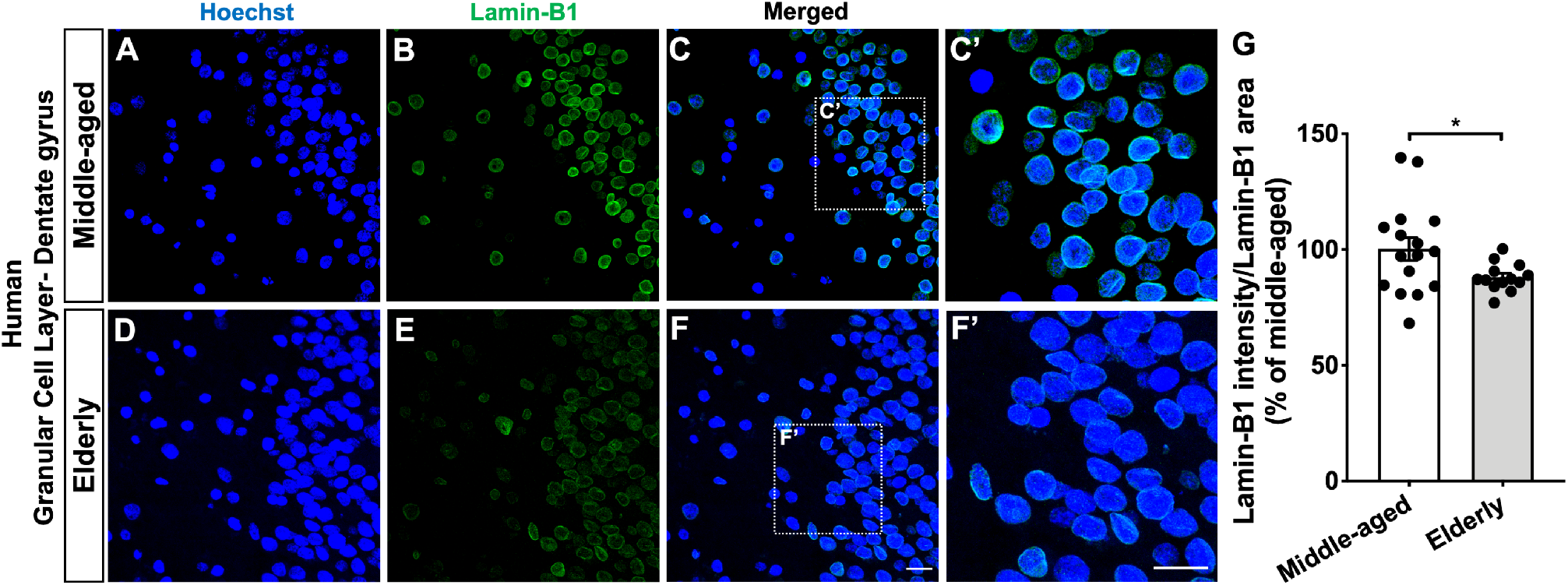
Lamin-B1 reduction in the human dentate gyrus upon aging. (A-G) Densitometric analysis of lamin-B1 staining at the hippocampal granular cell layer from *post-mortem* human tissue revealed an overall reduction of lamin-B1 intensity in elderly cases compared with middle-aged ones (p=0.0308). *n* = 16 and 13 individuals for middle-aged and elderly groups, respectively. Scale bars, 20 µm. Significance was determined using Unpaired t test with Welch’s correction. Error bar represent ± SEM. Individual data points are plotted and represent individual donors.

### Human hippocampal neural cells, including astrocytes, undergo nuclear abnormalities upon aging

Recent studies reported that lamin misregulation is associated with defects in neuronal nuclear morphology in age-related neurodegenerative diseases, including FTD and AD (Frost et al., 2016; Paonessa et al., 2019). However, it is still unknown whether these nuclear alterations also occur in neural cells during the human brain aging.

Thus, we first analyzed the percentage of regular, evaginated, invaginated and aberrant lamin-B1+ nuclei in the human hippocampal dentate gyrus from middle-aged and elderly donors. We observed a reduced proportion of regular nuclei at the hippocampal granular cell layer in elderly donors in comparison with middle-aged cases **(Figure 6 A-C)**, which was due to increased incidence of nuclear invaginations and aberrations **(Figure 6 A-C)**, accounting for a higher proportion of the total nuclear deformations in the elderly hippocampus compared with middle-aged cases **(Figure 6 D)**. We did not observe differences in the percentage of nuclear evaginations between middle-aged and elderly cases at the granular cell layer **(Figure 6 A-C)**. Therefore, these results indicate that neural cells, especially granule cells of the dentate gyrus, undergo nuclear invaginations and aberrations upon human aging.

**Figure 6.**
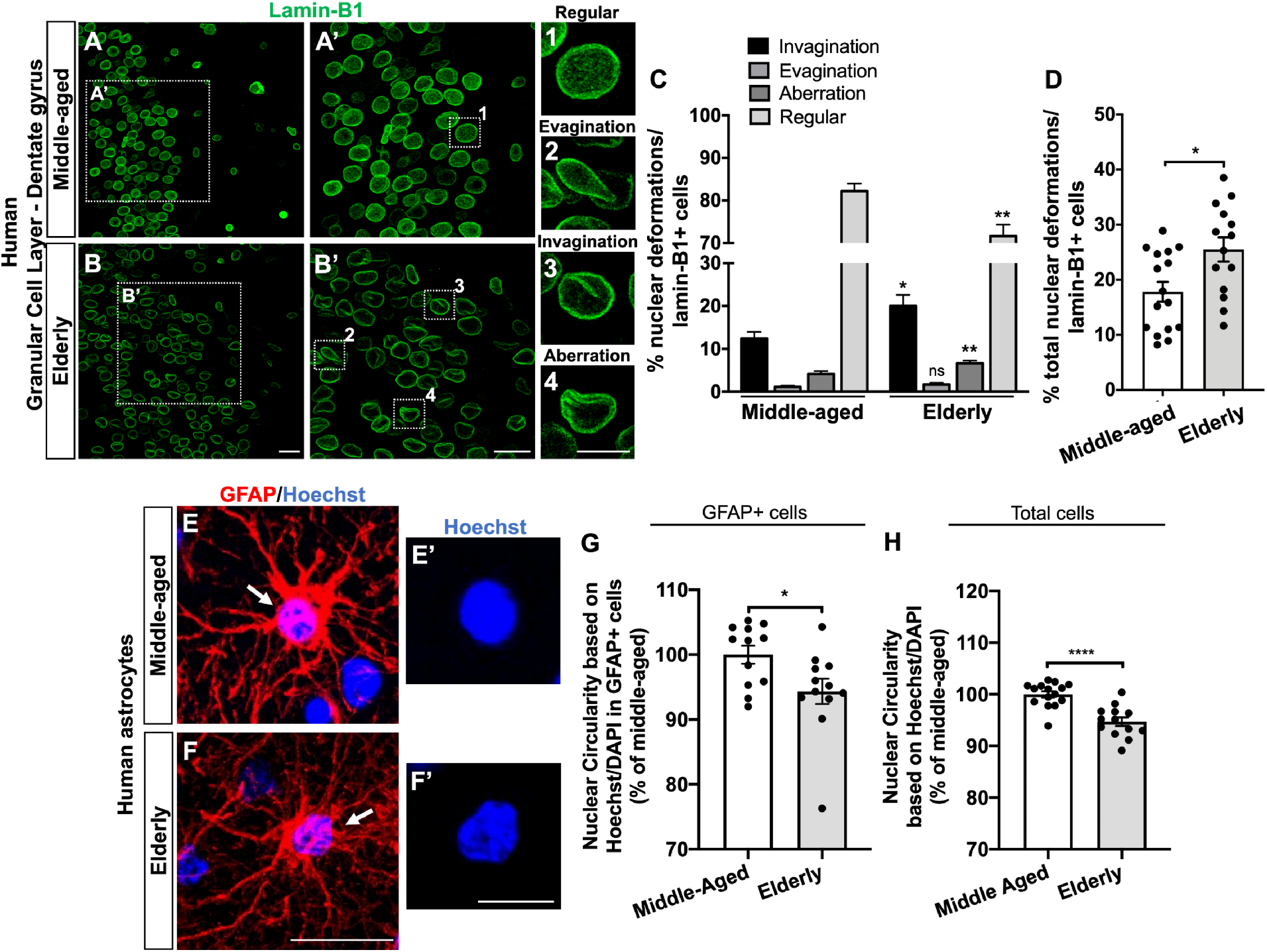
Neural cells, including astrocytes, from the granular cell layer of the human hippocampus undergo nuclear abnormalities upon aging. (A-B) Distinct nuclear morphological profiles were evaluated, such as regular (1), evagination (2), invagination (3) and aberration (4) at the hippocampal granular cell layer in *post-mortem* human tissue from middle-aged and elderly donors based on lamin-B1 staining. Scale bars, 20 µm in (B) and (B’); 10 µm in (4). (C) Elderly donors exhibited a higher incidence of invaginated (p=0.0494) and aberrant nuclei (p=0.0243), resulting in a decreased proportion of regular nuclei (p=0.0102), compared with the middle-aged group. *n*=16 and 14 individuals for middle-aged and elderly groups, respectively. (D) Elderly donors showed an increased proportion of total nuclear deformations (i.e., evagination + invagination + aberration) (p=0.0119). *n*=16 and 14 individuals for middle-aged and elderly groups, respectively. (E-G) Nuclear circularity was quantified based on Hoechst or DAPI staining in *post-mortem* human tissue from middle-aged (E-E’) and elderly donors (F-F’). Elderly donors presented a reduced nuclear circularity based on the number of GFAP+ cells (G; p=0.0275; *n*=12 individuals for both age groups) and on the total number of cells (H; p<0.0001; *n*=15 and 13 individuals for middle-aged and elderly groups, respectively). Scale bars, 20 µm in (F) and 10 µm in (F’). Significance was determined using Unpaired t test with Welch’s correction. Error bars represent ± SEM. Individual data points are plotted and represent individual donors.

Human astrocytes have several distinct features from their rodent counterparts including, morphological, molecular and functional properties (Barbar et al., 2020; Oberheim et al., 2009). We then asked whether nuclear abnormalities were also evident in the aged human astrocytes.

As we previously observed, cultured senescent astrocytes presented reduced levels of lamin-B1, which correlated with increased nuclear deformation and decreased nuclear circularity in these cells. Thus, to validate this data, we investigated the nuclear circularity of astrocytes of the granular cell layer from middle-aged and elderly cases. To do so, we used the Hoechst or DAPI nuclear staining, which provided a brighter signal of the nuclei boundaries of GFAP+ astrocytes in *post-mortem* human tissue. We observed a significant reduction of nuclear circularity in GFAP+ astrocytes from elderly donors compared with middle-aged ones **(Figure 6 E-G)**. Additionally, considering the total number of nuclei analyzed at the granular cell layer, we also reported a global reduction of nuclear circularity in elderly donors in comparison with middle-aged donors **(Figure 6 H)**.

Taken together, we conclude that the human aging is associated with defects in the nuclear morphology of neural cells, including astrocytes, and reduction of the levels of lamin-B1. Therefore, our data point to a new cellular and molecular marker to the human physiological brain aging.

## DISCUSSION

In the current study, we have identified that lamin-B1 loss and changes in nuclear morphology are hallmarks of astrocyte senescence *in vitro* and of neural cells, including astrocytes, in the aging mouse and human hippocampus. By using an *in vitro* model for astrocyte senescence, we observed that these cells undergo nuclear invaginations and changes in nuclear circularity, which were associated with the loss of lamin-B1. In agreement, these features were also observed in hippocampal astrocytes and granule cells in aged mice and in *post-mortem* human hippocampal tissue from non-demented elderly. Together, our data indicate that astrocytes undergo morphological and molecular modifications during aging and suggest that they may be important targets in understanding the basic mechanisms underlying human hippocampal aging as well as in identifying new age-associated biomarkers.

The hippocampus is critical for cognitive functions and is one of the most vulnerable brain regions to aging and implicated in age-related cognitive decline (Small et al., 2011). Moreover, emerging evidence has pointed to a differential vulnerability of hippocampal subfields in aging and neurodegenerative diseases (Fu et al., 2018; Morrison and Baxter, 2012; Roussarie et al., 2020), being the dentate gyrus particularly affected during normal aging (Morrison and Baxter, 2012; Nadal et al., 2020). Although most of these studies have focused on neuronal populations, recent findings have suggested that major changes in the gene expression profile of glial cells, especially astrocytes, might predict and/or impact the selective vulnerability of brain regions to aging-associated cognitive decline (Boisvert et al., 2018; Clarke et al., 2018; Soreq et al., 2017). Nevertheless, the contribution of astrocyte senescence to aging is far less studied.

The characterization of senescence-associated biomarkers is an important and firststep towards understanding the mechanisms underlying senescence itself and its role in the context of aging and age-related diseases. Emerging evidence shows that the loss of lamin-B1 is an important senescence-associated biomarker in several murine and human cell cultures/lines (Freund et al., 2012) and aging tissues, including in photoaged skin (Wang et al., 2017), thymus (Yue et al., 2019) lung (Saito et al., 2019), kidney and liver (Yousefzadeh et al., 2020). However, relatively less is known about the potential role of lamin-B1 in brain senescence. Here, we demonstrated that lamin-B1 loss is also a hallmark of astrocyte senescence *in vitro* and described for the first time that its reduction also occurs in hippocampal astrocytes from aged mice, as well as in the hippocampal dentate gyrus from elderly humans.

We first reported a global reduction of lamin-B1 in the hippocampal dentate gyrus from aged mice, which was partially due to a decreased proportion of hippocampal astrocytes expressing lamin-B1 in this age group. In agreement with our data, a recent study showed age-dependent downregulation of lamin-B1 in adult neural stem/progenitors cells in the mouse hippocampus, which underlies stem cell aging and negatively impacted adult neurogenesis (Bedrosian et al., 2020).

To further investigate lamin-B1 levels in astrocytes, we took advantage of an alternative long-term cell culture model for astrocyte senescence. Astrocytes cultured for 30-35 DIV recapitulated key features of the senescent phenotype, including increased SA-β-Gal activity and p16^INK4a^ expression, high expression of MMP3, IL-6, and reactive oxygen and nitrogen species, characteristics of SASP. These results corroborate other models previously described to study glial senescence (Bigagli et al., 2016, Pertusa et al., 2007; Bellaver et al., 2017; Souza et al., 2015) and reproduce several glial changes observed in the aging brain, including increased expression of p21, oxidative/nitrosative stress as well as an inflammatory phenotype (Bellaver et al., 2017). Accordingly, astrocytes acutely isolated from the cerebral cortex of aged mice showed a distinct gene expression signature, which also includes an increased inflammatory phenotype (Orre et al., 2014). Together, these results strongly support the use of long-term cell culture as an *in vitro* model to investigate the phenotype of glial cell senescence. As shown in murine hippocampus, we reported reduced levels of lamin-B1 in senescent cultured astrocytes associated with nuclear membrane malformations.

Human astrocytes are distinct from their murine counterparts, morphologically as well as molecularly, pharmacologically and, functionally (Barbar et al., 2020; Oberheim et al., 2009). Limited availability to human *post-mortem* samples has deeply hampered our understanding of brain aging at the cellular and molecular level. Whether the mechanisms underlying rodent brain aging apply to the human brain is still a matter of investigation. Here, we took the advantage of analyzing *post-mortem* human tissue from non-demented elderly and middle-aged donors. We observed reduced intensity of lamin-B1 in hippocampal cells at the granular cell layer from non-demented elderly donors. This was followed by increased nuclear deformations and a global reduction of nuclear circularity in the elderly hippocampus compared with middle-aged cases. Similarly to senescent mouse astrocytes *in vitro*, we observed a significant reduction of nuclear circularity in GFAP+ astrocytes from elderly donors compared with middle-aged ones, suggesting that alteration in nuclear properties might be a conserved characteristic of mice and human astrocyte aging. In agreement with our results, a recent *in vitro* study using X-irradiation induced senescent human astrocytes showed a downregulation of the lamin-B1 gene in these cells, together with higher expression of other classical senescent-associated biomarkers (Limbad et al., 2020).

Taken together, these findings strengthen the hypothesis that lamin-B1 loss is a hallmark associated with astrocyte senescence in mouse and human hippocampal aging. Although a direct link between lamin-B1 loss with changes in astrocyte phenotype and function has yet to be investigated, lamin-B1 downregulation has been recognized as a key step for progression of full cellular senescence in other cell types and tissues (Freund et al., 2012; Saito et al., 2019; Shimi et al., 2011), which can, in turn, contribute to tissue degeneration in aging and induce pathological conditions (Saito et al., 2019; Yue et al., 2019). Whether lamin-B1 loss is a consequence of the senescent phenotype or a trigger of astrocyte senescence remains to be investigated.

Lamin-B1 is a component of the nuclear lamina and is required for proper organogenesis, cell proliferation, self-maintenance, chromatin structure, nuclear integrity, and gene expression (Ho and Lammerding, 2012). Misregulation of nuclear lamins and abnormalities of the nuclear shape have been widely reported in laminopathies, including premature aging syndromes (Schreiber and Kennedy, 2013), and more recently in *in vivo* models for age-related neurodegenerative diseases. Reduction in lamin-B protein levels and nuclear invaginations have been shown in Tau-transgenic *Drosophila* brains, as well as in AD human brains, events associated with an aberrant cytoskeleton-nucleoskeleton coupling and neuronal death (Frost et al., 2016). Here, we report increased nuclear invagination and abnormal nuclear circularity in senescent astrocytes *in vitro* and in human aged brain. Likewise, a higher incidence of nuclear lamina invaginations is present in FTD human brain tissue and in FTD iPSC-derived neurons, which resulted in disrupted neuronal nucleocytoplasmic transport *in vitro* (Paonessa et al., 2019).

Astrocyte dysfunctions have been strongly implicated in the pathogenesis of several age-related diseases, although astrocyte senescence has only recently been fully addressed (Bussian et al., 2018; Chinta et al., 2018; Cohen and Torres, 2019). Recently, it has been shown that senescent astrocytes of PD patients were lamin-B1 deficient, while non-astrocytic neighboring cells retained basal levels of lamin-B1, suggesting that astrocytes may preferentially undergo senescence in PD brain tissue (Chinta et al., 2018). In this sense, this and other studies have shown that depletion of senescent cells, including senescent glial cells, mitigates PD pathology (Chinta et al., 2018), prevents tau-dependent pathology (Bussian et al., 2018), and cognitive deficits in AD mouse models (Zhang et al., 2019).

The fact that loss of lamin-B1 has been reported in age-related neurodegenerative diseases (Chinta et al., 2018; Frost et al., 2016; Paonessa et al., 2019) suggests that this feature may represent an early sign of age-associated brain pathology. Our study demonstrated that lamin-B1 downregulation and nuclear deformations are present in astrocyte (and other neural cells) from *post-mortem* human tissues from elderly donors, suggesting that astrocytes may play important role in the mechanisms underlying brain aging.

## EXPERIMENTAL PROCEDURES

### Animals

Newborn (P0) Swiss mice were used for astrocytes cultures. For *in vivo* experiments, we used male C57Bl/6 mice divided into two age groups: 2-3 months-old (young group) and 18-24 months old (aged group). All animals were housed at standard conditions with ad libitum access to food and water. Animal handling and experimental procedures were previously approved by the Animal Use Ethics Committee of the Federal University of Rio de Janeiro (CEUA-UFRJ, approval protocol 006/18) and the Central Authority for Scientific Experiments on Animals of the Netherlands (CCD, approval protocol AVD115002016659). Experiments were performed according to Brazilian Guidelines on Care and Use of Animals for Scientific and Teaching Purposes (DBCA) and the Directive of the European Parliament and of the Council of the European Union of 22 September 2010 (2010/63/EU).

### Human *post-mortem* brain material

Human *post-mortem* brain material was obtained from two brain banks to increase representation and statistical power: the Netherlands Brain Bank, Amsterdam (www.brainbank.nl) (NBB) and the Brain Bank of the Brazilian Aging Brain Study Group, University of São Paulo Medical School (http://en.gerolab.com.br/) (BBBABSG). A written informed consent for a brain autopsy and the use of the material and clinical information for research purposes had been obtained by the NBB (project number 1073) and BBBABSG (CAAE number: 30038520.0.0000.5257, approval number 3.986.070). We obtained paraffin-embedded tissue of the hippocampal area from both brain banks. A total of 30 donors were classified into two age groups: middle-aged (ranged from 50-60 years, n=16) and elderly (ranged from 76-93 years, n=14). Only non-demented controls were included in the analysis, based on the medical history and pathological scoring. Clinico-pathological information of all donors is present in **Supplementary Table 1**.

### Control and senescent astrocyte cultures

Primary cortical astrocyte cultures were derived from newborn Swiss mice as previously described (Matias et al., 2017) and the protocol for senescent astrocyte cultures was adapted from Bigagli et al., 2016. Briefly, cerebral cortices were removed, and the meninges were carefully stripped off. Tissues were maintained in Dulbecco’s minimum essential medium (DMEM) and nutrient mixture F12 (DMEM/F12, Invitrogen), supplemented with 10% fetal bovine serum (FBS, Invitrogen). Cultures were incubated at 37°C in a humidified 5% CO_2_, 95% air chamber for approximately 7 days *in vitro* (DIV) until confluence. Cultures were separated into two experimental groups: control and senescent astrocyte cultures. After confluence, control astrocyte cultures were treated with cytosine arabinoside (Ara C 10 μM, Sigma) in DMEM/F12 with 10% FBS for 48 h, washed and maintained in DMEM/F12 without FBS for additional 24 h until fixation or protein extraction. In contrast, after the treatment with Ara C, senescent astrocyte cultures were washed and maintained in DMEM/F12 supplemented with 10% FBS for 30-35 DIV, with medium exchange every 2 days. Twenty-four hours before fixation or protein extraction, senescent astrocyte cultures were incubated in DMEM/F12 without FBS.

### Immunocytochemistry of astrocyte cultures

Astrocyte cultures were fixed with 4% PFA in PBS (pH 7.4) for 15 min and nonspecific sites were blocked with 3% bovine serum albumin (BSA; Sigma-Aldrich), 5% normal goat serum (Sigma-Aldrich) and 0.2% Triton X-100 diluted in PBS for 1 h, before incubation with the following antibodies: rabbit anti-GFAP (1:1,000; DAKO Cytomation), rabbit anti-lamin-B1 (1:1,000; Abcam), rabbit anti-p16INK4a (1:100; Proteintech) and rabbit anti-iNOS (1:100; Abcam) at 4°C overnight. Subsequently, the cells were thoroughly washed with PBS and incubated with secondary antibodies at room temperature (RT) for 2 h. Secondary antibodies were Alexa Fluor 546-conjugated goat anti-rabbit IgG or goat anti-mouse IgG (1:1,000; Invitrogen), or Alexa Fluor 488-conjugated goat anti-rabbit IgG or goat anti-mouse IgG (1:300; Invitrogen). Nuclei were counterstained with DAPI (Sigma-Aldrich) and cells were observed with a TE2000 Nikon microscope.

### Immunohistochemistry of mouse brain tissue

The animals were euthanized by i.p. injection of a lethal dose of sodium pentobarbital (Euthanimal; Alfasan BV, Woerden, The Netherlands) and then transcardially perfused with PBS. Brains were removed and fixed with paraformaldehyde 4% at 4°C for 24 h, then kept in PBS at 4°C. Forty-µm thick sagittal sections were obtained using vibratome (Leica) and subjected to immunohistochemistry. Sections were incubated with blocking buffer composed of 5% normal donkey serum (NDS, Sigma-Aldrich), 2% BSA (Sigma-Aldrich), 1% Triton X-100 diluted in PBS for 1 h, before incubation with the following primary antibodies: rabbit anti-lamin-B1 (1:1,000; Abcam) and mouse anti-GFAP (1:1,000; Sigma-Aldrich) at 4°C for 24 h. The slices were then washed with PBS and incubated with secondary antibodies at RT for 2 h. Secondary antibodies were Alexa Fluor 488-conjugated donkey anti-rabbit IgG (1:1,400; Jackson Immuno Research Inc.) or Cy3-conjugated donkey anti-mouse IgG (1:1,400; Jackson Immuno Research Inc.). Nuclei were counterstained with Hoechst 33528 (1,1:000; Thermo Fisher Scientific) and coverslips were mounted in Mowiol (0.1 M Tris, pH 8.5, 25% glycerol, 10% w/v Mowiol 4-88 [Sigma-Aldrich]). Sections were imaged on a confocal microscope Zeiss LSM 880.

### Immunohistochemistry of human paraffin-embedded hippocampal tissue

Immunohistochemistry of human paraffin-embedded tissue was done according to a modified protocol (van Strien et al., 2014). Paraffin sections (7 µm thick) were deparaffinized, rehydrated, and washed in distilled water, followed by PBS/0.05% Tween 20 for 30 min. Hereafter, sections were incubated with PBS/0.3% hydrogen peroxide for 10 min, followed by antigen retrieval through exposure heating in a steamer in citrate buffer (10 mM citric acid, 0.05% Tween 20, pH 6.0; 98°C) for 20 min. After cooling down to RT, nonspecific sites were blocked with 5% NDS (Sigma-Aldrich), 2% BSA (Sigma-Aldrich), 0.1% Triton X-100 diluted in PBS for 1 h, before incubation with the following primary antibodies: rabbit anti-lamin-B1 (1:1,000; Abcam), mouse anti-GFAP (1:1,000; Sigma-Aldrich) or mouse anti-GFAP (1:1,000; Millipore) at 4°C overnight. The sections were then washed in PBS and incubated with secondary antibodies at RT for 2 h. Secondary antibodies were Alexa Fluor 488-conjugated donkey anti-rabbit IgG (1:1,400; Jackson Immuno Research Inc.) or Cy3-conjugated donkey anti-mouse IgG (1:1,400; Jackson Immuno Research Inc.). Next, sections were washed in PBS and incubated in Sudan Black solution (0.3% Sudan Black in 70% ethanol) for 7 min to quench autofluorescence, and then washed in 70% ethanol for 1 min, followed by an additional wash in PBS. Nuclei were counterstained with Hoechst 33528 or DAPI and coverslips were mounted in Mowiol (Sigma-Aldrich). Sections were imaged on a confocal microscope (Zeiss LSM 880 or Leica TCS SPE).

### SA-β-Galactosidase (SA-β-Gal) activity

The SA-β-Gal activity was performed with the SA-β-Gal staining kit (Cell Signaling) according to the manufacturer’s instructions. Briefly, control and senescent astrocyte cultures were incubated with the fixative solution at RT for 15 min, washed with PBS, and incubated with the β-galactosidase staining solution at 37°C in a dry incubator overnight. Subsequently, the SA-β-Gal staining was quantified under a TE2000 Nikon microscope. The optical density of the X-Gal color was determined with the Threshold color plugin from Image J software (NIH, USA), which converts the blue color to optical density. The optical density of each field was normalized by the number of cells labeled by DAPI.

### Reactive oxygen species measurement

Dihydroethidium (DHE; Invitrogen) was freshly prepared before each experiment. Control and senescent astrocytes cultures were loaded with DHE at a final concentration of 10 µM for 40 min, and cells were immediately imaged on a TE2000 Nikon microscope. DHE intensity was quantified using ImageJ software as previously described (Diniz et al., 2017).

### Nitrite measurement

Nitric oxide (NO) production was determined indirectly through the assay of nitrite (NO_2_), a stable metabolite of NO, according to the Griess reaction (Ding et al., 1998). Briefly, a 50 µL aliquot of control or senescent astrocyte conditioned medium was mixed with an equal volume of Griess reagent [0.1% N-(1-naphthyl) ethylenediamine dihydrochloride, 1% sulfanilamide, and 2.5% phosphoric acid] and incubated at 22°C for 10 min, followed by the absorbance measurement at 540 nm. Based on a standard curve of NaNO_2_ (Sigma-Aldrich) ranging from 0 to 100 µM, nitrite concentration was calculated. Background NO_2_^-^was subtracted from each experimental value.

### Quantitative RT-PCR (qPCR)

The cortical astrocytes were lysed with TRIzol® (Invitrogen), and total RNA was isolated and purified with Direct-zol™ MiniPrep Plus (Zymo Research, Irvine, CA, USA) according to the manufacturer’s protocol. The RNA was quantified using a Nano Drop ND-1000 spectrophotometer (Thermo Fisher Scientific, Waltham, MA, USA). The total RNA (1-2 μg) was reverse transcribed with a GoScript™ Reverse Transcriptase cDNA reverse transcription kit according to the manufacturer’s instructions (Promega Corporation, an affiliate of Promega Biotecnologia do Brasil, Ltda). Primers were designed and synthesized by IDT-DNA (San Diego, CA, USA). The specific forward and reverse oligonucleotides were as follows: p16^INK4a^: (F) CAG CTC TTC TGC TCA ACT AC, (R) CGC ACG ATG TCT TGA TGT; MMP3: (F) CTG AAG GAG AGG CTG ACA TA, (R) GAG CAG CAA CCA GGA ATA G; IL-1β: (F) CAG GCA GGC AGT ATC ACT CA, (R) TAA TGG GAA CGT CAC ACA CC; IL-6: (F) CAA AGC CAG AGT CCT TCA GAG, (R) TGG TCC TTA GCC ACT CCT TC; and the reference gene RPLP0: (F) CAG GTG TTT GAC AAC GGC AGC ATT, (R) ACT CAG TCT CCA CAG ACA ATG CCA. Quantitative real-time PCR was performed using Fast SYBR Green Master Mix qPCR Master Mix (Applied Biosystem ™); the cycling conditions were 95°C for 20 sec, and 40 cycles of 95°C for 1 sec, 60°C for 20 sec in the Quant Studio 7 Flex System (Applied Biosystem ™). The relative expression levels of the genes were calculated using the 2−ΔΔCT method (Livak and Schmittgen, 2001).

### Western blotting

Protein concentration in cell extracts was measured by the BCA Protein Assay Kit (Cole-Parmer). Forty micrograms protein/lane was electrophoretically separated on a 10% SDS polyacrylamide gel and electrically transferred onto a Hybond-P PVDF transfer membrane (Millipore) for 1.5 h. Membranes were blocked in PBS-milk 5% at RT for 1 h. Next, membranes were incubated in block solution overnight with the following primary antibodies: rabbit anti-lamin-B1 (1:1,000; Abcam) and mouse anti-GAPDH (1:1,000; Abcam). Membranes were incubated for 1 h with IRDye 680CW goat anti-mouse antibody and IRDye 800CW goat anti-rabbit antibody (LI-COR, 1:20,000) and then scanned and analyzed using Un-Scan-It gel version 6.1 (Silk Scientific).

### Densitometric analysis of mouse tissue

Hippocampal dentate gyrus, comprising the granular cell layer and molecular layer was imaged on a confocal microscope (Zeiss LSM 880). Densitometry for the immunohistochemistry images was performed using integrated density values generated by the Fiji software (NIH, USA), and represent the mean of 2 brain tissue sections per mouse, with 2-3 images per section. Results represent the mean of four animals per experimental group, as indicated in the graph and legend.

### Densitometric analysis of astrocyte cultures

Densitometry for the immunocytochemistry images was performed using integrated density values generated by the ImageJ software (NIH, USA), and normalized by the number of cells per field. At least 10-15 images were acquired from duplicate coverslips per experimental condition. For each result, the exact number of astrocyte cultures per experimental group is indicated in the graph and legend.

### Nuclear deformations and circularity measurements on astrocyte cultures

Control and senescent astrocyte cultures were immunostained for lamin-B1, counterstained with DAPI and images were acquired with a TE2000 Nikon microscope with a 63x objective. Based on the previous classification of nuclear deformations through super-resolution fluorescence microscopy and 3D reconstruction, we quantified the relative number of nuclear deformations based on the total number of lamin-B1+ nuclei per image. A total of 3,084 cells from 9 control astrocyte cultures and 4,481 cells from 10 senescent astrocyte cultures were analyzed. We also calculated the nuclear circularity per lamin-B1+ nuclei by using the Fiji software and by employing the formula: circularity = 4π (area/perimeter^2^). Circularity has a maximum value of 1 and diminishes as the nuclear shape becomes increasingly convoluted (Zhang et al., 2018). A total of 1,231 and 944 lamin-B1+ nuclei, respectively, from 7 and 6, control and senescent astrocyte cultures, were analyzed. Astrocytes cultures were prepared from different litters.

### Densitometric analysis of human paraffin-embedded tissue

Human paraffin-embedded hippocampal tissues were immunostained for lamin-B1 and counterstained with Hoechst or DAPI. The granular cell layer and polymorphic layer of the hippocampal dentate gyrus were imaged on a confocal microscope (Zeiss LSM 880 or Leica TCS SPE), using the same image parameters for middle-aged and elderly groups. Densitometry for the immunohistochemistry images was performed using the values of integrated density and nuclear area generated with the Fiji software (NIH, USA). The integrated density was normalized by the area of lamin-B1 immunostaining. The values represent the mean of 2-4 independent hippocampal tissue sections per donor. For each result, the exact number of donors are indicated in the graph and legend.

### Nuclear deformations and circularity measurements of human paraffin-embedded tissue

Human paraffin-embedded hippocampal tissues were immunostained for lamin-B1 and GFAP, and counterstained with Hoechst or DAPI. We used the same parameters described above (for astrocyte cultures) to quantify the number of invaginated, evaginated, and aberrant nuclei based on lamin-B1 staining. The relative number of nuclear deformations was expressed based on the total number of lamin-B1+ nuclei per image. Tissue sections were imaged on a confocal microscope (Zeiss LSM 880 or Leica TCS SPE), using the same image parameters for middle-aged and elderly groups. Nuclear circularity was calculated per Hoechst (tissue from the NBB) or DAPI (tissue from the BBBABSG) labelled nuclei, as well as in GFAP+ cells at the granular cell layer from middle-aged and elderly donors. The total number of cells and nuclei analyzed for each marker and experimental condition is described in **Supplementary Table 2**.

### Statistical analysis

Statistical analysis was done by Student’s t-test, using GraphPad Prism version 8 (GraphPad Software, La Jolla, CA, USA). P-value <0.05 was considered statistically significant. Error bars represent the standard error of the mean (SEM). For each result, the exact number of experiments, animal samples, or human *post-mortem* hippocampal tissue donors are indicated in the graph, legend, and the respective “Experimental procedures” section.

## ACKNOWLEDGEMENTS

We thank Marcelo Meloni, Grasiela Ventura, Jacqueline Sluijs, Daniëlle Vonk and Roland van Dijk for technical assistance. This work was supported by grants from Conselho Nacional de Desenvolvimento Científico e Tecnológico (CNPq) (IM, IVD, LSN, FCAG), Coordenação de Aperfeiçoamento de Pessoal de Nível Superior (CAPES) (IM, LPD), Fundação Carlos Chagas Filho de Amparo à Pesquisa do Estado do Rio de Janeiro (FAPERJ) (IVD, FCAG), Fundação de Amparo à Pesquisa do Estado de São Paulo (FAPESP) (REPL, CS, RN, WJF, LTG), U.S. Department of Health & Human Services, National Institute of Health (LTG), ZonMW Memorabel and Alzheimer Nederland (EMH, JM), Departamento de Ciência e Tecnologia, Ministério da Saúde (Decit-MS) (IM, LPD, APBA, GV, FCAG), Instituto Nacional de Neurociência Translacional (INCT-INNT) (FCAG).

## CONFLICT OF INTEREST

The authors declare no conflict of interest.

## AUTHOR CONTRIBUTIONS

IM, EMH, FCAG conceptualization; IM, LPD, IVD, EMH, JM, and FCAG research design; IM, LPD, IVD, LSN, APBA, GV research performance; IM, LPD, IVD, LSN, APBA, GV, data analysis; EMH, JM, FCAG supervision; IM writing-original draft preparation; IM and FCAG reviewing and editing; REPL, CKS, RN, WJF, LTG; data analysis and resources; EMH, JM, FCAG funding acquisition and project administration.

## DATA AVAILABILITY STATEMENT

The data that support the findings of this study are available from the corresponding author upon reasonable request.

## Abbreviations

AD: Alzheimer’s disease
Ara C: cytosine arabinoside
BSA: bovine serum albumin
CNS: central nervous system
DHE: dihydroethidium
DIV: days *in vitro*
DMEM: dulbecco’s minimum essential medium
FBS: fetal bovine serum
FTD: Frontotemporal Dementia
GAPDH: glyceraldehyde 3-phosphate dehydrogenase
GFAP: glial fibrillary acidic protein
iNOS: inducible nitric oxide synthase
MMP3: metalloproteinase 3
NO: nitric oxide
NO_2_^-^: nitrite
PBS: phosphate-buffered saline
PD: Parkinson’s disease
SASP: senescence-associated secretory phenotype
SA-β-Gal: senescence-associated β-galactosidase activity.

## Notes

### Competing Interest Statement

The authors have declared no competing interest.

