## Supplemental Tables for "Loss of lamin-B1 and defective nuclear morphology are hallmarks of astrocyte senescence *in vitro* and in the aging human hippocampus"

**Matias et al., 2021**
**Supplementary Tables**

**Supplementary Table 1. Donors.**

| Sample | Sex | Age | Diagnosis | Amyloid | Braak | ApoE | PMD | pH CSF | Cause of death | Source |
|---|---|---|---|---|---|---|---|---|---|---|
| 1991-118 | m | 59 | Non-demented control | - | 0 | 33 | <06:00 | 6.05 | Aspiration pneumonia and thrombo-emboli | NBB |
| 1996-014 | f | 54 | Non-demented control | 0 | 0 | 33 | 08:00 | 6.45 | Acute renal failure | NBB |
| 1998-125 | f | 58 | Non-demented control | - | 1 | 43 | 06:15 | 6.31 | Multiple organ failure | NBB |
| 2005-034 | m | 56 | Non-demented control | 0 | 0 | 33 | 14:00 | 7.03 | Terminal congestive heart failure | NBB |
| 2011-081 | m | 55 | Non-demented control | 0 | 0 | 33 | 07:30 | 6.88 | *Euthanasia with esophageal cancer | NBB |
| 2012-071 | f | 57 | Non-demented control | - | 0 | 32 | 07:40 | 6.47 | *Euthanasia with metastatic urothelial cancer | NBB |
| FLA24 | f | 60 | Non-demented control | 0 | 1 | - | 11:18 | - | Pulmonary embolism | BBBABSG |
| FLA9 | m | 56 | Non-demented control | 0 | 2 | 34 | 18:00 | 6.00 | Heart failure | BBBABSG |
| FLA3 | f | 56 | Non-demented control | A | 1 | 33 | 13:00 | 7.00 | Cirrhosis | BBBABSG |
| FLA6 | f | 53 | Non-demented control | - | - | - | 21:42 | 6.80 | Heart faliure | BBBABSG |
| FLA21 | f | 50 | Non-demented control | 0 | I | - | 10:36 | - | Pneumonia | BBBABSG |
| FLA12 | f | 56 | Non-demented control | 0 | 0 | - | 16:24 | - | Myocardial infarction | BBBABSG |
| FLA15 | f | 54 | Non-demented control | 0 | 2 | 33 | 09:12 | - | Pulmonary edema | BBBABSG |
| FLA18 | f | 57 | Non-demented control | 0 | 2 | - | 11:18 | - | Myocardial infarction | BBBABSG |
| FLA30 | f | 59 | Non-demented control | - | - | - | 15:06 | 7.00 | Acute pancreatitis | BBBABSG |
| FLA27 | m | 60 | Non-demented control | 0 | 0 | 33 | 12:15 | 6.90 | Heart failure | BBBABSG |
| 2000-007 | m | 85 | Non-demented control | 0 | 2 | 33 | 15:10 | 6.85 | Myocard infarction | NBB |
| 2005-020 | m | 79 | Non-demented control | A | 1 | 33 | 06:30 | 6.32 | Cerebrovascular accident | NBB |
| 2005-061 | f | 93 | Non-demented control | 0 | 2 | 33 | 05:50 | - | Cachexia (mamma carcinoma) | NBB |
| 2006-049 | f | 84 | Non-demented control | 0 | 1 | 33 | 04:45 | 6.26 | Heart failure, lung emphysema and dehydration | NBB |
| 2008-054 | f | 92 | Non-demented control | A | 1 | - | 07:00 | 6.55 | Acute death, probably pulmonary emboly | NBB |
| 2011-028 | f | 81 | Non-demented control | 0 | 1 | 33 | 04:25 | 6.67 | Intestinal ischemia | NBB |
| 2011-111 | m | 93 | Non-demented control | 0 | 1 | 33 | 05:05 | - | Heart failure | NBB |

| | | | | | | | | | | |
|---|---|---|---|---|---|---|---|---|---|---|
| 2013-016 | m | 83 | Non-demented control | A | 1 | 33 | 05:15 | 6.60 | Myocardial infarction and palliative sedation | NBB |
| FLA11 | f | 76 | Non-demented control | A | 1 | 33 | 15:12 | 6.74 | Heart failure | BBBABSG |
| FLA8 | m | 84 | Non-demented control | 0 | 2 | 23 | 17:48 | 6.74 | Heart failure | BBBABSG |
| FLA5 | f | 79 | Non-demented control | 0 | 2 | 33 | 11:36 | 6.17 | Pulmonary thromboembolism | BBBABSG |
| FLA14 | f | 79 | Non-demented control | 0 | 2 | - | 16:48 | 6.20 | Pulmonary edema | BBBABSG |
| FLA20 | f | 77 | Non-demented control | A | 2 | 34 | 11:18 | 6.50 | Heart failure | BBBABSG |
| FLA23 | f | 91 | Non-demented control | A | 2 | 33 | 17:00 | - | Coronary artery disease | BBBABSG |

NBB = Netherlands Brain Bank number; BBBABSG: Brain Bank of the Brazilian Aging Brain Study Group; m = male, f = female; ApoE = Apolipoprotein E; PMD = *Post-mortem* Delay in hours: min; CSF: Cerebrospinal fluid. *Euthanasia is legal in the Netherlands.

**Supplementary Table 2.** Number of nuclei analyzed for the nuclear deformation and circularity measurements in *post-mortem* human hippocampal tissue

| Marker | Measurement | Middle-aged | | Elderly | |
|---|---|---|---|---|---|
| | | GCL | Nº of donors | GCL | Nº of donors |
| Lamin-B1+ nuclei | Nuclear deformation | 4,191 | 16 | 4,522 | 14 |
| Hoechst/DAPI nuclei | Nuclear circularity | 741 | 15 | 715 | 13 |
| Hoechst/DAPI of GFAP+ cells | Nuclear circularity | 156 | 14 | 136 | 12 |
